# Dynamic walking behavior during odor trail-following in locusts

**DOI:** 10.1101/2023.02.22.529569

**Authors:** Mike Traner, Barani Raman

## Abstract

One of the important subsets of odor sources used in olfactory navigation is surface-bound sources, which can broadly take the form of point sources or trails. Odor trails, in particular, have been observed to be highly relevant components of olfactory-based navigation for species as broadly distributed as dogs and ants. Here, we present an automated treadmill setup capable of dynamically printing odor trails of arbitrary lengths and configurations, and with closed-loop control of speed based on the subject’s movement. We used this setup to characterize trail-following behavior in locusts (*Schistocerca americana*). The free-moving behavior of the locusts is more naturalistic and is richer in plumbable data than many traditional assays. We reveal broad classes of behavioral walking motifs and their dynamic transitions as locusts pursue or avoid an odor trail. Furthermore, we show how these motifs vary across individuals, with the identity of the odorant and with respect to the sex of the organism. Our dataset and analyses provide a first demonstration that this model organism is capable of robust odor trail following, and provides a comprehensive analysis of the dynamic motifs that underlie this behavioral capability.

## Introduction

Odor trails have been observed to be highly relevant components of olfactory-based navigation for species as broadly distributed as dogs^1–4^ and ants^5–7^. However, despite their ecological prevalence, few studies have examined animal behavior around odor trails compared to airborne plume tracking during flight. Among the important work that has been done on the subject are studies on the extraordinary capabilities of large animals such as dogs, wolves, rattlesnakes, and other predators to track the odor trails left by prey over long distances^1,8–12^. Smaller animals, in particular mammals such as rats and mice^13^, and invertebrates such as ants^5–7,14^ have also been successfully used to examine trail-following behavior.

The capabilities of many moths and other insects^15–18^ to follow the odor plumes produced by conspecifics are common examples of the capabilities of olfaction-based navigation. While odor plume tracking displays many similarities to surface-bound trail following, many fundamental differences remain.

Particularly, in odor plume tracking the fluid nature of the medium results in greater distortion and dispersal of the odorant due to air currents, which drive sharp drops in concentration as the distance from the source grows. Unlike odor plume tracking, during odor trail following the animals stay close to and in most cases on the odor source, with constant exposure to the tracked odorant. Irrespective of the type of odor-driven navigation, a number of important characteristics have been found across species^5,13,19–21^. Such commonalities include the sinusoidal swings back and forth during trail and plume following^5,13^ and casting-like behavior upon losing the scent^13,19^. However, surface-bound odorants are also distinct from airborne odorants in that disturbing the bound medium can increase the concentration of the odorant available to detect^22–24^, which may drive sniffing^13^ and antennal sampling strategies^5^ when exploring odor trails.

Compared to natural trails, artificial trails used to study trail-following in a lab suffer from a few key limitations. Most studies utilizing small animals in a laboratory setting were performed in a fixed arena, where one or more odor trails were applied prior to the experiment, either by pipetting odorant onto the arena surface prior to placement of the animal or by releasing animals to form their own scent trails before performing manipulations ^5,7,25^. The former approach was used to characterize antennal sampling patterns in ants while actively tracking an odor trail^5^. The drawback of this approach arises from the size of the arena and therefore the limited trail length, and the inability to alter the pattern of the odor trail during an experiment. To overcome some of these limitations, a looped treadmill where the trail had been predefined and pipetted onto the track prior to introducing the animal has been used to study navigation strategies. This approach was used to evaluate the head movements of rats following a reward-conditioned odor trail, where Khan et al. were able to evaluate the capability of rats to track trails while monitoring respiration, nose movements, and olfactory bulb LFPs^13^. They were able to demonstrate that rats can track even with one nostril closed, as well as model the tracking behavior as the rat followed, and lost, the trail. This setup had the distinct advantage that while the animal’s movement was still constrained in two dimensions, the subject was able to explore an unlimited distance in the third dimension (i.e. the length of the trail), which was a key advantage given the large size of rats. However, the pre-pipetting of the odorant locked in the configuration of the trail for the entirety of each trial. Perhaps of equal importance, each revolution of the treadmill would reintroduce any residual odorant left by the animal, which could alter its behavior.

We developed an odor trail virtual reality system designed to alleviate some of the shortcomings in previously utilized experimental setups. We demonstrate capabilities to dynamically print single-use odor trails onto paper towels that pass through the behavior chamber only once, produce arbitrary trail configurations, permit unlimited exploration in one direction, and control movement of the treadmill via closed-loop control driven by the movement of the animal. We use this approach to characterize trail-following behavior in locusts (*Schistocerca americana*). The free-moving behavior of the locusts is both more naturalistic and produces richer, more plumbable data than many traditional assays applied to locusts, e.g. palp opening response^26^ or t-maze assay^27^. We utilize this rich repository of behavior to further examine broad classes of behavior and how they vary with odorant condition, sex, and time.

## Methods

### Treadmill Design

The Odor Trail VR (virtual reality) system is designed around a core behavior chamber where the animal subject is placed to interact with the trail, and a surrounding set of supporting modules mounted on an 80-20 aluminum extrusion frame. There are numerous challenges to studying trail-following behavior. One of the most important is constraints on the size of arenas; the larger the model organism, the larger the necessary arena, which has likely contributed to why many previous studies have focused on ants^5,7^. However, Khan et al. addressed this constraint with the innovation of using a treadmill design^13^, which enables exploration in one direction for an infinite distance while maintaining a small arena size. Constraining the size of the arena has other advantages as well, such as allowing detailed recording of the entire arena with a single camera.

In order to leverage the same advantages as Khan et al., the Odor Trail VR was designed around a treadmill design concept. However, a number of problems are presented when dealing with other organisms, and the design sought to address these while also resolving some shortcomings of Khan et al.,’s original treadmill. One constraint of the original design, at least in trials where a looped belt was used, is the possibility of an animal encountering residues of its own scent when the belt loops over. This could influence behavior as the scent residue gradually accumulates with each loop, and is particularly likely in animals such as ants where many species lay pheromone trails as they navigate^7^. This is resolved by exclusively utilizing an open-belt design, where the belt is discarded after the animal passes over it once. A white paper towel (Scott, Number 12388) was selected for the treadmill ‘belt,’ and was mounted and tensioned at the front of the treadmill, with alignment and tensioning achieved using custom mechanisms.

Another constraint of the design utilized by Khan et al. was that the treadmill was operated at a continuous speed. The continuous speed is unsuitable for animal models that have sporadic periods of movement such as locusts because the animal would be dragged against the back end of the arena which could disturb their natural behavior. To prevent this, a closed-loop control system was implemented, with the centroid position of the locust determining the speed at which the treadmill operated. A webcam (Logitech C920) was mounted above the behavior chamber, and a python script was implemented to track the position of the locust within the chamber. When the locust was in the rear half of the chamber the speed would be set to zero, and as the locust passed the midline the speed would exponentially increase to a maximum (2.6 cm/s) as it approached the front of the chamber. The movement of the belt was controlled by a DC motor linked to a paper drive assembly from an inkjet printer (HP Photosmart C6180). The movement was monitored using the built-in quadrature encoder and the speed was regulated using a PID (Proportional Integral Derivative) control system; this ensured smooth and continuous movement, minimizing the risk of disturbing the locust when changing speed.

One other potential issue comes from the pre-trial pipetting of the odor trail; during the course of the trial, the odor trail may dissipate. Pausing the treadmill during stationary periods of the animal would exacerbate this issue. To alleviate this potential confound, the odorant was dynamically printed onto the treadmill prior to the belt arriving at the behavior chamber. To enable the largest range of odorants, a pressurized odorant system was selected for the printing method; odorants were dissolved at a defined concentration in a dichloromethane solution, then pressurized and released via a solenoid (Clippard NR3-2-12) using timed pulses through brass 0.1mm printheads (M6 Brass 3D Printer printheads, Shenzhen Aopin Network Technology Co., LTD). Under visible light conditions, all odorant solutions were mixed with alcohol dyes to produce a uniform color (see Odorant Solution). The printhead was constructed by mating the pressurized delivery system printhead to a print carriage assembly from the same HP inkjet printer (HP PN C9087-60014 Rev A2). The position of the printhead was regulated by a second PID system, with the set position controlled by an equation using the treadmill belt position as input. By setting the sequence of equations and transition points between them, a range of arbitrary trail patterns could be dynamically produced on the belt prior to entry into the behavioral chamber. Once printed, the paper towel passed through a solvent evaporation tunnel, where the dichloromethane was exhausted and the trail dried prior to reaching the behavior chamber.

Finally, the behavior chamber itself was designed to enable minimally-inhibited exploration by the locust and easy tracking from the overlooking camera. To encourage moving forward on the treadmill, the negative gravitaxis displayed by locusts and many other organisms was taken advantage of by inclining the treadmill at 15 degrees. By also coating the walls with Fluon (Fluon, byFormica; a PTFE suspension that when dried, makes climbing difficult for arthropods) the locusts were more likely to ascend towards the front of the behavior chamber. Furthermore, the chamber was underlit by a pair of diffused LED arrays that could be independently switched on; for visible light white LEDs were switched on, while for dark conditions IR LEDs were used. Under both conditions, the backlighting enabled the consistent acquisition of the locust position in the chamber by the python script. A glass lid over the top of the chamber ensured that jumping locusts were not able to escape the arena.

Electronics were incorporated into a custom PCB. Sensor reading and motor outputs for the treadmill were handled by a Teensy 3.5 microcontroller (Teensy 3.5, PJRC), in order to take advantage of the two built-in hardware quadrature decoders. Print carriage position was first established using a limit switch, with the position monitored thereafter using an optical encoder strip. Both the print carriage and the treadmill belt driver utilized the Avago quadrature encoders and optical gratings retrieved from the inkjet printer. Odorant print pulses and motor PWM output were controlled using TIP31C transistors.

The python controlling and data-logging program communicated with the Teensy 3.5 using a serial connection. Timestamped position sensor data was transmitted to the python program, which in turn would send the designated speed for the treadmill to the microcontroller based on the position of the locust in the behavior chamber. Printhead position equations would be sent, either in the form of the coefficients of a linear equation with the slope, offset, and start position specified, or as the amplitude, phase offset, and start position of a sinusoidal equation. All images retrieved from the camera were in parallel processed to find the locust centroid, timestamped using the computer clock, compressed to jpeg format, and saved to multipage tiff files for later analysis. A new tiff file was generated every 200 frames, due to tests finding that the time to write the image into the file increased beyond that point on the experimental computer. Serial messages from the microcontroller were also timestamped with the computer time and saved, as was the recorded position of the locust in each frame. The timing was compared and all events synchronized to a common time frame for analysis.

### Odorant Solution

The odorant solution was prepared by mixing 1% v/v pure odorant compounds in dichloromethane for all odorants except frass extract. Frass extract was prepared by drying frass under a vacuum for 24 hours, soaking the dried frass in dichloromethane at a 1:1 ratio of DCM to dried frass for an additional 24 hours, and then decanting and vacuum-filtering the liquid using Schleicher and Schuell grade 576 circular filters. In order to maintain consistency across the panel, all odorant solutions were dyed using Pinata brand (1147 Healdsburg Ave, Healdsburg, California, 95448) alcohol dyes, at a ratio of 2.5mL Burro Brown, 2mL Sunbright Yellow, and 1mL Rainforest Green to 150mL DCM to dye the solution a uniform brown shade matching the frass extract. Odorant solutions were used within two weeks of preparation, or upon the formation of solids at the upper edge of the solution, whichever came first.

### Locust Selection

Post fifth-instar American locusts (*Schistocerca americana*) from a crowded colony were used for all experiments.

### Trial Structure

Recordings lasted 15 minutes, during which locusts were allowed to roam freely in the behavior chamber. Prior to the beginning of each trial, the treadmill was manually advanced so that a fresh odor trail was present in the behavior chamber. The locust was then deposited on the middle of the odor trail at the rear of the chamber, the chamber lid closed, and recording began. Locusts were not constrained following release and prior to the beginning of the recording and therefore were able to move away from the start point prior to the start of recording. After 15 minutes elapsed, the recording was stopped, the locust was retrieved and placed into a labeled cup for storage, and the treadmill re-primed for the next trial.

### Experimental Design & Data Collection

The first application of the treadmill was to evaluate whether trail-following behavior occurred in locusts and to quantify exploratory behavior around surface-bound odor trails. While early testing data was collected using more complex ‘zig-zag’ trails, it was found that locusts tended to exhibit an attraction to the walls. When the trail approached the walls, locusts were highly likely to leave the trail and walk beside the wall as they continued their ascent. In order to minimize this confound and simplify analyses, all trails in the experiment consisted of straight lines along the middle of the behavioral chamber unless otherwise indicated. Similarly, locusts were found to move significantly more under visible-light conditions, so unless otherwise indicated all trials were collected under visible-light conditions with the trail dyed to a uniform brown color.

Experiments were structured to use two sequences to control for sequence effects, one with the odor trail first, and the other with the control trail first. Daily trials were structured by first dividing the daily lot of locusts into two batches. All trials would be preceded by a priming phase where the belt was run with the behavior chamber empty to ensure a fresh trail was present during the beginning of the trial. The first batch of locusts would be placed in the behavior chamber with a control trail, where they would be recorded for 15 minutes. They would then be removed and set aside in an isolated cup. Once all locusts in the first batch had been run on the control trial, the control solution was flushed and the treadmill was re-primed with the odorant solution for the day. The locusts were run again in sequence, with each being recorded for 15 minutes with the odorant trail. After completion of the first batch under the odor condition, the second batch of locusts would be recorded under the odor condition first, with each locust being set aside after their trial. Once the second batch had been recorded, the odorant solution was flushed from the print system, and the treadmill was re-primed with the control solution again. The second batch of locusts was then run sequentially under the control condition. All batches were sought to be roughly balanced in the sex ratio of the locusts, to ensure that representative samples were obtained for all odorants used. However, due to mistaken identification during initial locust retrieval or early cessation of experiments, some odorants display imbalances in the sex ratios of locusts.

A panel of 20 odorants at a 1% v/v concentration in the odorant solution was assayed, with an additional 4 odorants assayed at both 1% v/v and 10% v/v concentrations.

### Tortuosity Analysis

Tortuosity was measured by taking the ratio of the total of the 0.1s sampled trajectory length to the total displacement during the trial. This was then averaged across all locusts for each condition.

### Trajectory Analysis

Position data for each trial was uniformly resampled at 10 Hz for all analyses. Position data in pixels was subsequently transformed into cm for the relevant analyses, such as distance traveled and velocity analyses. Data was assembled for locusts across all odorants and conditions. The vector of each 0.1s sample was computed, and the sum of all vector lengths was then summed for the 15-minute trial period to find the total distance traversed. The trajectory was then segmented along its length into 50 pixel (1.05 cm) segments for behavioral classification.

Locust movement segments were classified into four movement categories: stationary periods, walking in straight lines (the type of movement with the lowest tortuosity), tortuous walking (curved, turning, and other low-displacement high-distance movements), and circling behavior (a special case of maximally tortuous walking). The angle between each point and the subsequent point (the angle of the trajectory segment) was found relative to the positive y-axis (straight forward on the treadmill). The change in the angle of each segment relative to the locust was then found by defining the first point as 0, and each subsequent point as the change in angle relative to the previous point.

Segments where more than 60 seconds elapsed without any movement were defined as stationary periods and excluded from further classification. Segments were classified as circling behavior when the locust consistently turned in one direction such that at least 360 degrees of rotation was achieved. If one segment was not classified as circling but was bounded on both sides by three continuous sections of circling, then it was defined as circling for the purposes of bridging the sections and accounting for perturbations caused by, e.g. encountering a wall. Segments were then classified as straight walking if at least three consecutive sections were within +/-15 degrees of straight, such that all consecutive segments totaled to a cumulative angle change of fewer than 45 degrees. Finally, all remaining segments of movement were classified as tortuous movement.

The proportion of locusts in each of the four states over time was computed by averaging the proportion of locusts classified for each category in each time sample (1/10^th^ of a second). The significance of trends over time was computed using Spearman’s correlation of the proportion of a category over time, averaged across locusts, with significance evaluated using 10,000 permutations. Comparisons between category classification over time were computed by bootstrapping the difference between the Spearman’s rho values using 20,000 bootstraps, and evaluating whether the 0.05 confidence interval excluded 0.

### Trail Following Analysis

To demonstrate the utility of the treadmill as an exploratory behavior assay, we collected behavior for sixteen odorants at a single concentration, and for four additional odorants at two different concentrations. We then evaluated the tendency of locusts to remain in proximity to the trail (attraction index). Locusts were recorded for 15 minutes in both a control trial (where the trail solution consisted only of dichloromethane and dye) and an odor trial (where the trail solution also contained either 1% or 10% v/v odorant). The sequence of the presentations and sex distribution was divided roughly equally.

A proximity index was determined by first resampling the position of the locust centroid at ten samples/second. The distance of the locust to the trail at each time point was then divided by the maximum distance possible from the trail, that is the distance between the center of the trail to the arena edge (**Fig. 3b**). The trail proximity index (TPI) was then calculated by subtracting the ratio from one, and averaging across all time points. The resulting number then ranges from 0 to 1, where a number close to 1 indicated the locust stayed on the trail the entire time, while a number close to 0 indicated the locust spent the entire time as far from the trail as possible.

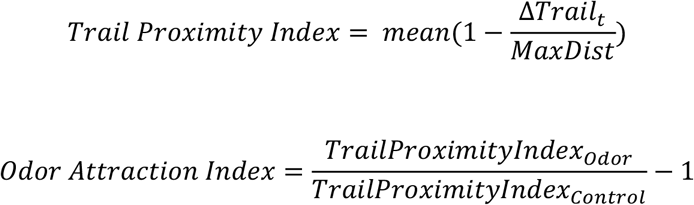

Finally, the mean ratio was taken across all locusts for each odorant using both metrics, and the significance of the difference between the odor and control trial was evaluated for significance using a Wilcoxon Signed-Rank Test at p<0.05 (**Fig 3a**).

### Markov Modelling of Locomotion

Prior to computing the transition probabilities, the sampling rate had to be computed, due to the time-sensitivity of Markov models. Bouts of each of the four categories were defined, and the median of all bout durations was then used as the sampling rate for determining states for Markov transition probabilities.

The categorization of each locust was sampled at the median bout duration for the 15-minute trial, then the transition matrix was computed for each locust. The transition matrices were averaged across locusts, and the differences between subsets (control and odor, male and female, etc.) were computed for comparison.

### Comparison of correlations over time

Correlations over time were computed using Spearman’s non-parametric rho. A p=0.001 confidence interval of the difference between the correlations in the conditions being compared was computed using 20,000 bootstraps and declared a significant difference if the confidence interval excluded zero.

### Data Availability

Data and code used for generation of all figures and results will be made publicly available in an online open access repository (figshare.com) up on acceptance of the manuscript. In addition, the design files and rig software and firmware will also be made available in a public database (github).

### Other Methodology and Statistics

- A Monte Carlo simulation was performed to evaluate the stability of the Trail Proximity Index (TPI) values. 100 samples were drawn with replacement from each odorant, and the mean correlation with the overall TPI was evaluated. For a sample size of n ≥ 15 the R^2^ correlation value exceeded 0.7. Therefore, n ≥ 15 locusts was used for all odorants.
- The following exclusion criteria were used:
  ○ Any trial missing data due to technical malfunctions.
  ○ Any trial where the locust was able to hang on the glass behavioral chamber lid, thereby avoiding interaction with the treadmill surface.
- Locusts were gathered each day after feeding time, and after dividing by sex, each locust was randomly assigned to the control-first or odorant-first condition trial groups.
- All data was quantified using automated algorithms.
- All statistical tests performed were non-parametric to minimize assumptions about the data distributions unless otherwise specified.

## Results

### Olfactory VR Treadmill for studying odor-driven navigation

We began by designing an olfactory treadmill that can be adapted in a closed-loop fashion based on the behavior of each individual locust (**Fig. 1a, b, Supp. Fig 1a-c**). The length of each trail was not predefined but was a function of the amount of walking observed in each animal and therefore varied across individuals. The paper towels on which the odor trail was printed and that formed the walking substrate of the treadmill passed the behavioral arena only once, thereby limiting potential confounds that arise from animal subject residues. After the odor solution was printed onto the paper towel, it passed through a solvent-exhaust tunnel where it was dried prior to entering the behavioral chamber. The chamber was back-lit with either visible or IR-LEDs to allow tracking of the locust movement. Together with the capability to print odor trails of arbitrary shapes (**Fig. 1c-g**), the designed olfactory treadmill addressed the key challenges previously noted for studying surface-bound olfactory navigation.

**Figure 1:**
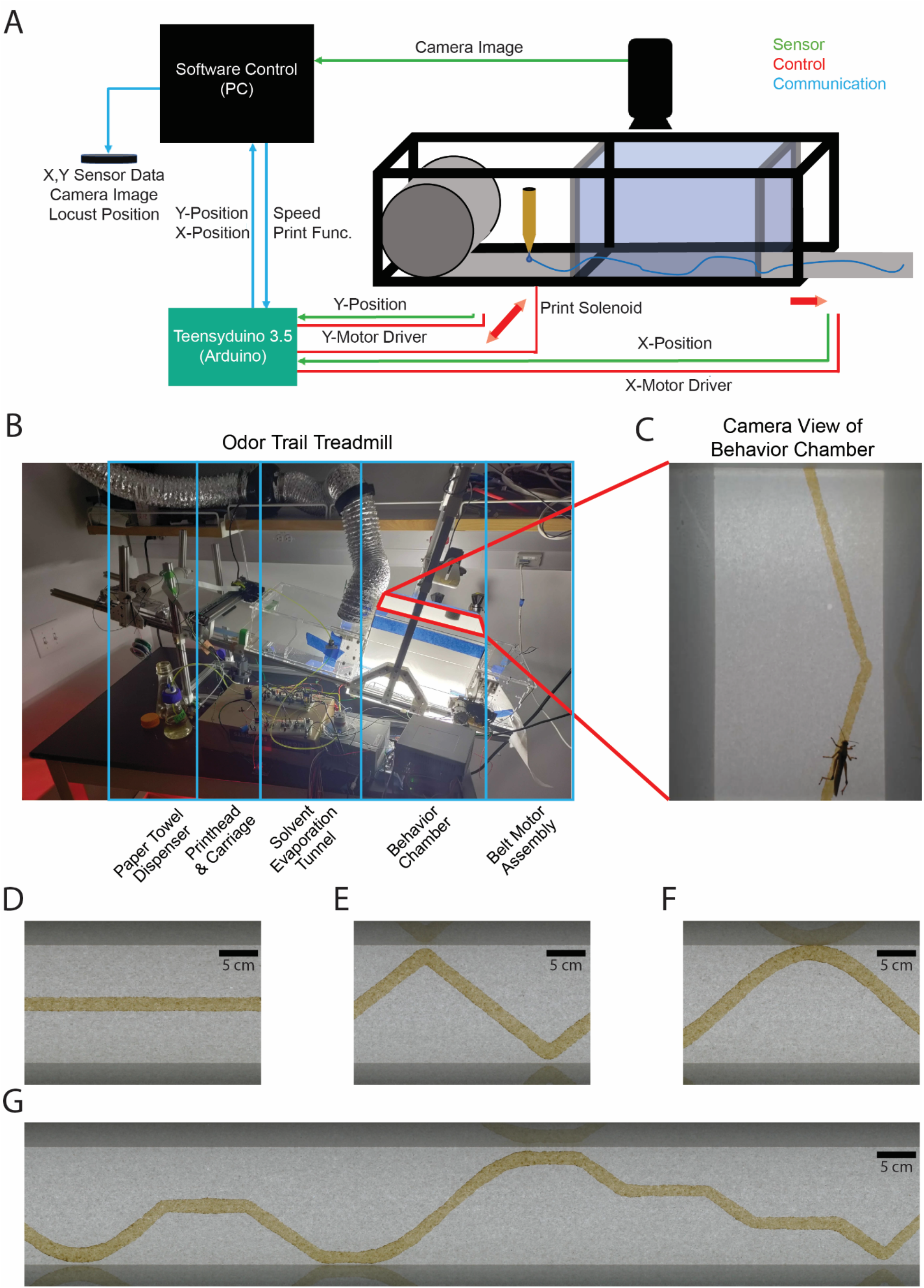
An automated odor trail treadmill for tracking locust behavior. **A)** A schematic of the treadmill system with various components and information flow between them (indicated by lines) is shown. Sensors signals used for control and data logging are indicated in green, and control signals, i.e. outputs to drive motors or solenoids, are indicated in red. The sensor and control signals assembled are stored on disk for subsequent analyses. **B)** A photo of the built treadmill with the 4 primary modules highlighted is shown. The first module is a dispenser for the paper towel on which the trail is printed. The second module is the print mechanism, with the printhead consisting of a nozzle that releases the odorant solution onto the paper towel from a pressurized reservoir when a solenoid opens. The printhead is mounted to the print carriage, which can move back and forth to enable the printing of arbitrary patterns. The third module is the solvent evaporation tunnel, where the solvent to prepare odor solutions is allowed to evaporate and exhaust. The fourth module is the behavior chamber, where the locust interacts with the dried trail. A camera mounted above the behavioral chamber records the locust position, and the chamber is underlit by either IR or visible lights. The fifth and last module is the belt motor assembly, where the paper towel is pulled via a geared motor, and after exiting is deposited on the floor for disposal. **C)** Top view of the behavior chamber (module 3) from the view of the camera mounted above the chamber is shown. The position of the locust is monitored in order to enable closed-loop control of the treadmill: When the locust is in the bottom (rear) half of the chamber, the treadmill does not move. However, when the locust advances past the midline towards the top (front) of the chamber, the speed is exponentially increased until the maximum speed of 2.6 cm/s is reached. Odor trails are dyed a uniform brown to match the color of the locust frass extract odorant (see Methods). **D-G)** Odor trails of different patterns that were printed using the printhead (module 2) are shown. Note that a straight line parallel to the length of the chamber at a defined offset was the most basic pattern that was used for the majority of our experiments. Panel G shows a complex pattern formed by mixing a sequence of straight, straight-sloped, and sinusoidal lines.

### Locusts can follow trails left by surface-bound odorants

Can locusts follow odor trails? While several chemicals have been identified as putative pheromones and aggregation cues^28–34^, it remains to be shown whether locusts can use olfactory cues for the purposes of navigation. We began to examine this issue with the automated olfactory treadmill. We used zig-zag and straight trails of geraniol (the primary component of rose oil) to determine whether or not locusts could track this odorant. Geraniol was chosen as the trail odorant because we have previously found that locusts are attracted by this odorant in a T-maze assay (Saha et al, 2023).

Representative traces of locust walking through the treadmill are overlaid atop a control trail (solvent without any odorant; blue) and a 10% Geraniol odorant trail (red trace; **Fig. 2a-e**). We found that locusts do engage in trail-following behavior. This was observed irrespective of whether the trail was straight or zig-zagged (**Fig. 2a-e, Supp. Fig 2a-c**). To eliminate the potential confound due to the presence of the visual cues, we repeated the experiments under visible (**Fig. 2a-c**) and IR (**Fig 2d, e**) illumination. We found that locusts engage in trail following in both conditions, although differences in the tortuosity were noted between the visible (mean tortuosity: odor = 1.26e5, control = 0.82e5; see Methods) and IR (tortuosity: odor = 0.40e5, control = 0.25e5). In the remainder of the study, we used straight odor trails in visible conditions for simplicity.

**Figure 2:**
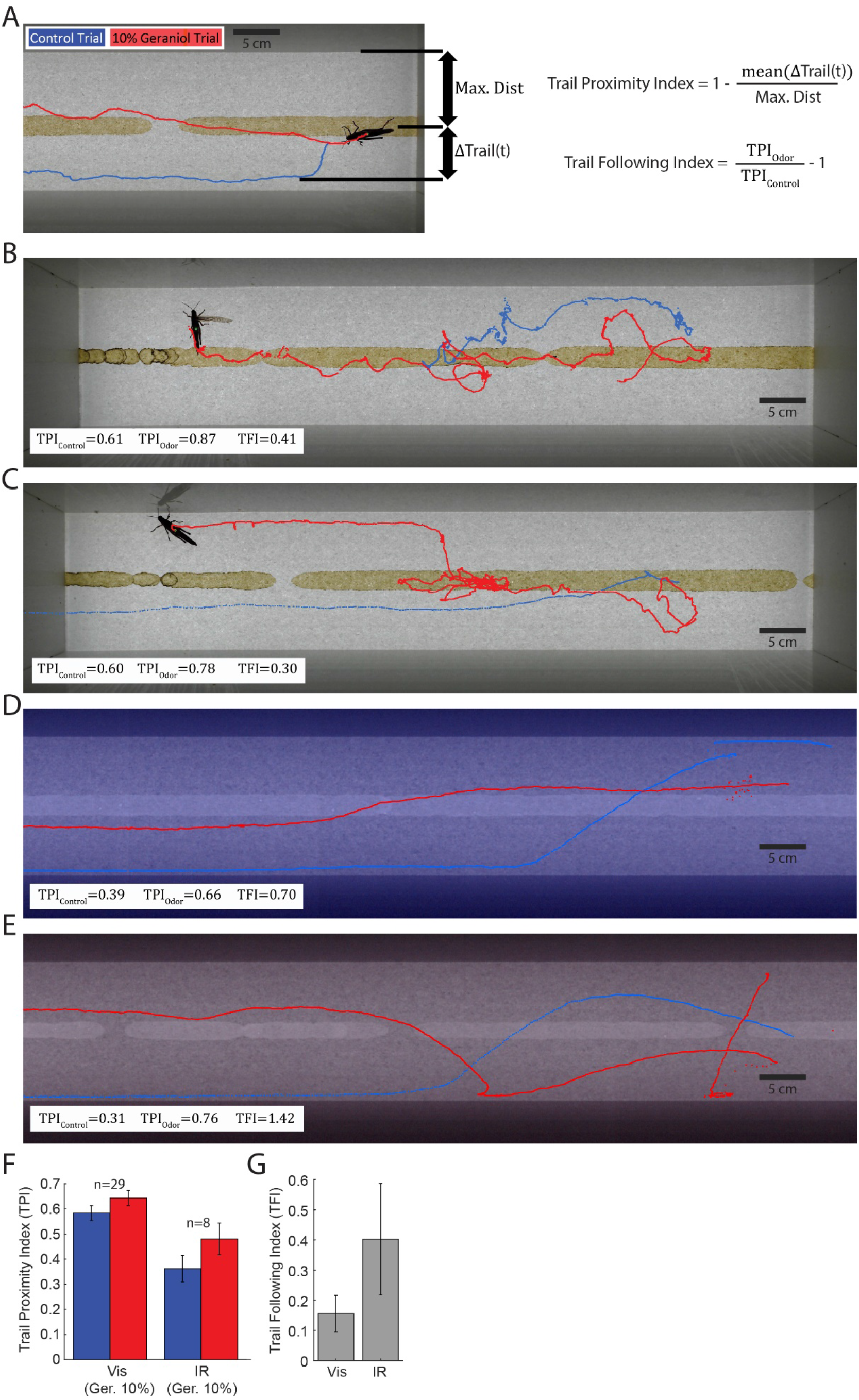
Locusts can follow an odor trail. **A)** Locust walking trajectories exhibited by a single locust while exploring a 10% geraniol trail (in red) or control trial (solvent in blue) are shown. The location of the straight-line geraniol trail that was printed onto the walking substrate is also shown for reference. A similar straight-line trail was also used for the control condition (not shown). Two different indices were computed based on the locust walking patterns: *Trail Proximity Index (TPI) and Trail Following Index (TFI)*. TPI is a measure of how much time the locust spent near the trail on a single trial. A TPI value of 0 indicates the locust avoided the trail the entire time, and a TPI value of 1 indicates the locust spent the whole time on the trail. Trail Following Index measures how close the locust was to the trail during the odor trial compared to the control trial. A negative TFI value indicates the locust spent much less time by the odor trail relative to the control trail, and a positive value that the locust spend more time close to the odor trail than what the same locust did in the control trail. **B, C)** Example locust trajectories of two different locusts under visible conditions. The TPI values for odor and control trails and the TFI values were computed for each locust and shown in the plots. **D, E)** Example trajectories for two different locusts under IR conditions. Again, the TPI values for odor and control trails and the TFI values were computed for each locust and shown in the plots. **F)** Quantification of TPI for odor and control conditions under visible and IR conditions. Locusts spent more time near the geraniol 10% trail under both visible and IR conditions. **G)** Quantification of Trail Following Index for visible and IR conditions. Locusts spent more time near the geraniol trail under both conditions. (n = 29 and 8 for visible and IR conditions respectively, same as for **Panel F**)

**Figure 3:**
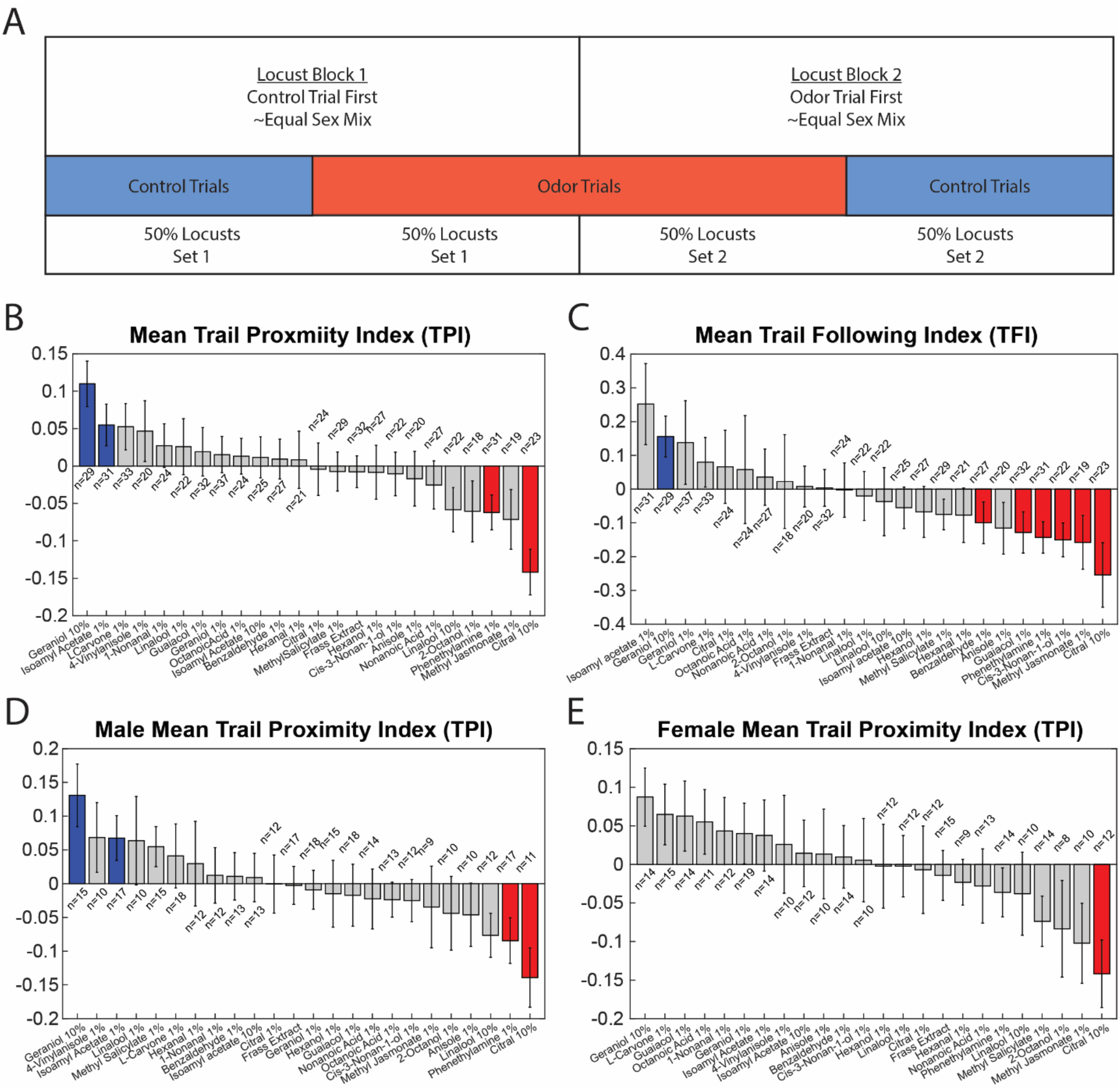
Odor trail following preferences in locusts. **A)** Experiment structure for trail following assays. Locusts for the day were first divided into two groups, split as evenly as possible. The first group of locusts was run on the control trials first, with the second group run on the odor trials first. This structure was repeated each day of the experiment. The exploration behavior of a total of 619 locusts was tracked using this experimental structure. **B)** The mean trail proximity index (TPI) for twenty-four odorants (twenty odorants at 1% concentration and four odorants at 1% and 10% concentration) are shown. Odorants significantly above the median of all odorants are indicated in blue (Wilcoxon Signed-Rank Test at p<0.05), while those significantly below the median are indicated in red. Geraniol 10% and Isoamyl Acetate (IAA) 1% were significantly attractive, while Citral 10% and Phenethylamine 1% were significantly repellant. **C)** The mean trail following index (TFI) for the same twenty-four odorants in panel A were computed and shown here as a bar plot. The ordering of odorant based on TFI was very similar to the TPI except for a few odorants such as Isoamyl Acetate 1% (significantly different TPI between the odor and control trails; Wilcoxon Signed-Rank Test p<0.05). Multiple odorants were significantly repellant using the TFI metric, including Citral at 10%, Methyl Jasmonate at 1%, cis-3-Nonan-1-ol at 1%, Phenethylamine at 1%, Guaiacol at 1%, and Benzaldehyde at 1%. A large number of repellant odors become significant suggesting that the test may be more sensitive to repellant odors than attractive odors. **D)** TPI computed for twenty-four odorants but looking only at male locusts: the same odorants were significant as for the overall TPI, namely Geraniol 10% and Isoamyl Acetate 1% were significantly attractive, and Citral 10% and Phenethylamine 1% were significantly repellant. **E)** TPI computed for only the female locusts: only one odorant was significantly different from the median, Citral 10%, which was significantly repellant.

To quantify locust trail-following behavior, we used two metrics, the Trail Proximity Index (TPI), the ratio between time spent near the trail and time spent away from the trail, and the Trail Following Index (TFI), which is the zero-centered ratio of the odor and control condition TPIs (**Fig 2a**, see Methods for details). As can be noted, locusts spent more time near the geraniol trials compared to the control case (**Fig. 2f; the** trail of the dyed solvent without odorant).

In addition, to account for variations across individuals, each locust was run on both odorant and control trials (**Fig. 2h**). The Trail Proximity Index of the odor and control condition was then used to compute an adjusted Trail Following Index (**Fig. 2a**, see Methods for details) which accounts for attraction to or repellency from the trail by factors other than the odorant. Using this metric, we similarly find a trail-following preference for geraniol 10% over control (**Fig. 2g**).

In sum, these results reveal that like ants^5^ and dogs^1,2,2^ locusts do have the capability to track trails left behind by surface-bound odorants.

### Trail-following vs. trail-avoiding preferences

Next, we wondered if the trail-following preference varies depending on the identity of the odorant. To examine this we used a diverse odorant panel that included green leaf volatiles, putative pheromones, and aggregation cues. In total, we tracked the behavioral preference of 610 locusts (319 males and 291 females). For each locust, we tracked the odor trail following preference for both the odor trail and the control trail (**Fig**. **2a, h**).

We found that trail-following preference varied a great deal across the odor panel. Note that some odorants like geraniol and isoamyl acetate induced positive trail following preference (i.e. more time spent near the odor trail; **Fig. 3b**). Locusts spent more time away from the trail when odorants such as citral and phenethylamine were used, indicating the repulsive nature of these odorants. Significant deviation from the overall median response, and from the control odor trials are identified using colored bars (**Fig. 3a, c, d;** two-sided Wilcoxon signed rank test p<0.05, **Fig 3b**, paired two-sided Wilcoxon signed rank test p<0.05).

### Dynamics of locomotory behavior

Next, we examined the distribution of velocities during the first, second, and third 5-minute segments of the experimental trial. We found that the locust velocity varied over time, with a larger proportion of higher-velocity segments during the later parts of the trial (**Fig. 4a**). We also found that the control condition developed a ‘hump’ in the velocity distribution around 2cm/s (**Fig. 4b**), suggesting that movement patterns are time, as well as odor condition dependent.

**Figure 4:**
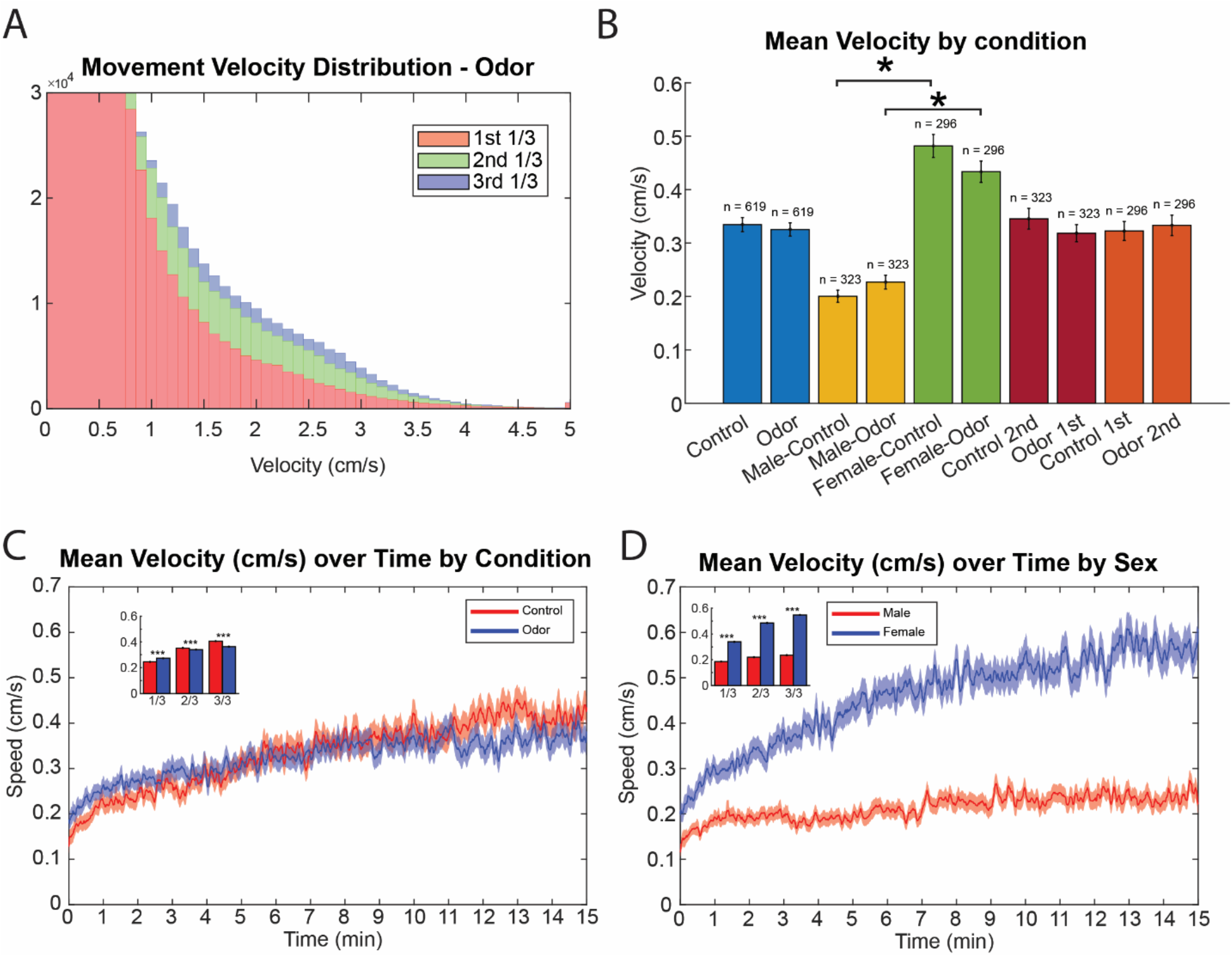
Locust walking speed varies over time. **A)** The distribution of walking velocities under the odor condition is shown for all locusts (n = 619). The walking velocity distribution for the initial five minutes (1/3^rd^) of the trial is indicated in red, the middle five minutes in green, and the last 5 minutes in blue (each trial – 15 minutes in duration). The walking speed became greater during the later portions of the trail. As a result, the green and blue distribution of velocities had a right-tail that included higher velocity values. **B)** A comparison of the average walking velocities between the odor and control conditions overall is shown. Comparisons are also made between walking velocities of male and female locusts in odor and control conditions, and to examine whether or not the presentation order mattered. The difference between overall mean walking speed between odor and control was not significant in any subset, however, the differences in walking speeds between males and females were dramatic and significant (asterisk indicates significance level of p < 0.05, Wilcoxon Rank-Sum test. Error bars indicate s.e.m.) **C)** Walking velocity as a function of time for the control and odor conditions are shown (n = 619). While walking speed in odor conditions slowly increases, it largely plateaus around the 8-minute mark. In contrast, the walking speed in the control condition continues to increase throughout the 15-minute period. The differences in the distribution of velocities for odor and control conditions at each 0.1s sample for each 5-minute subset are quantified in the inset plot, and all are significant. This suggests that the presence of an odor spurs an initial amount of exploration (i.e. increased walking speed), which falls off, while the control condition begins more slowly but gradually rises. (Envelopes indicate s.e.m.) **D)** Walking velocity as a function of time for male and female locusts during combined odor and control conditions (male n = 323, female n = 296). Consistent with **panel C**, females move much faster than males. Both males and females continue to move faster during the course of the trial, however, females have a much higher rate of increase during the trial. (Envelopes indicate s.e.m.)

We then examined locust mean velocity under the odor and control conditions (**Fig. 4c**, for details, see Methods). The velocity was similar regardless of whether the trail was an odor trail or the control trail. Further, the sequence of exposure, the odorant trail first or control trail first, did not significantly alter the average velocity. However, the amount of distance walked varied significantly between the male and female locusts (p<0.001 for both conditions; two-sided Wilcoxon signed-rank test).

We then further examined velocity as a function of time (**Fig 4d, e**). In general, consistent with the results shown in **Fig. 4a**, the walking velocities increased in the later time segments of an experimental trial. We found that walking velocities in odor condition was greater than those observed during the control conditions only during the early time segments and that the trend switched after 7-8 minutes (**Fig. 4d**). In contrast with the male walking speed, the walking velocity in female locusts was much more elevated and remained higher while exploring odor trails than the control trails throughout the experimental trial (**Fig. 4e**). This result suggests that the exploration/walking strategies are dynamic and differ between male and female locusts.

### Locomotory behavioral motifs and their dynamics in odor-driven navigation

To further understand how locusts followed odor trails, we segmented movement trajectories into one of four categories or exploration motifs: stationary periods of non-movement, linear exploration or straight walking, curved or tortuous walking, and circling where there is complete rotation, thereby capturing non-movement as well as the extremes (circling and straight walking) and intermediates (curved walking) of tortuosity (**Fig. 5a**; see Methods). As can be noted, all movement categories can be observed in most locust trajectories (**Fig. 5a**). The distribution of each of these motifs during both odor and control trial explorations is shown for both male and female locusts in **Fig. 5b, c**. As can be noted, trajectories for both odor and control conditions were characterized by roughly equal proportions of stationary periods, straight walking, and tortuous walking, with considerably less circling behavior. However, in the odor condition, significantly less time was spent stationary, and more time was spent in tortuous walking, than in the control condition (p=0.017, p=0.005 respectively; two-sided Wilcoxon signed-rank test).

**Figure 5:**
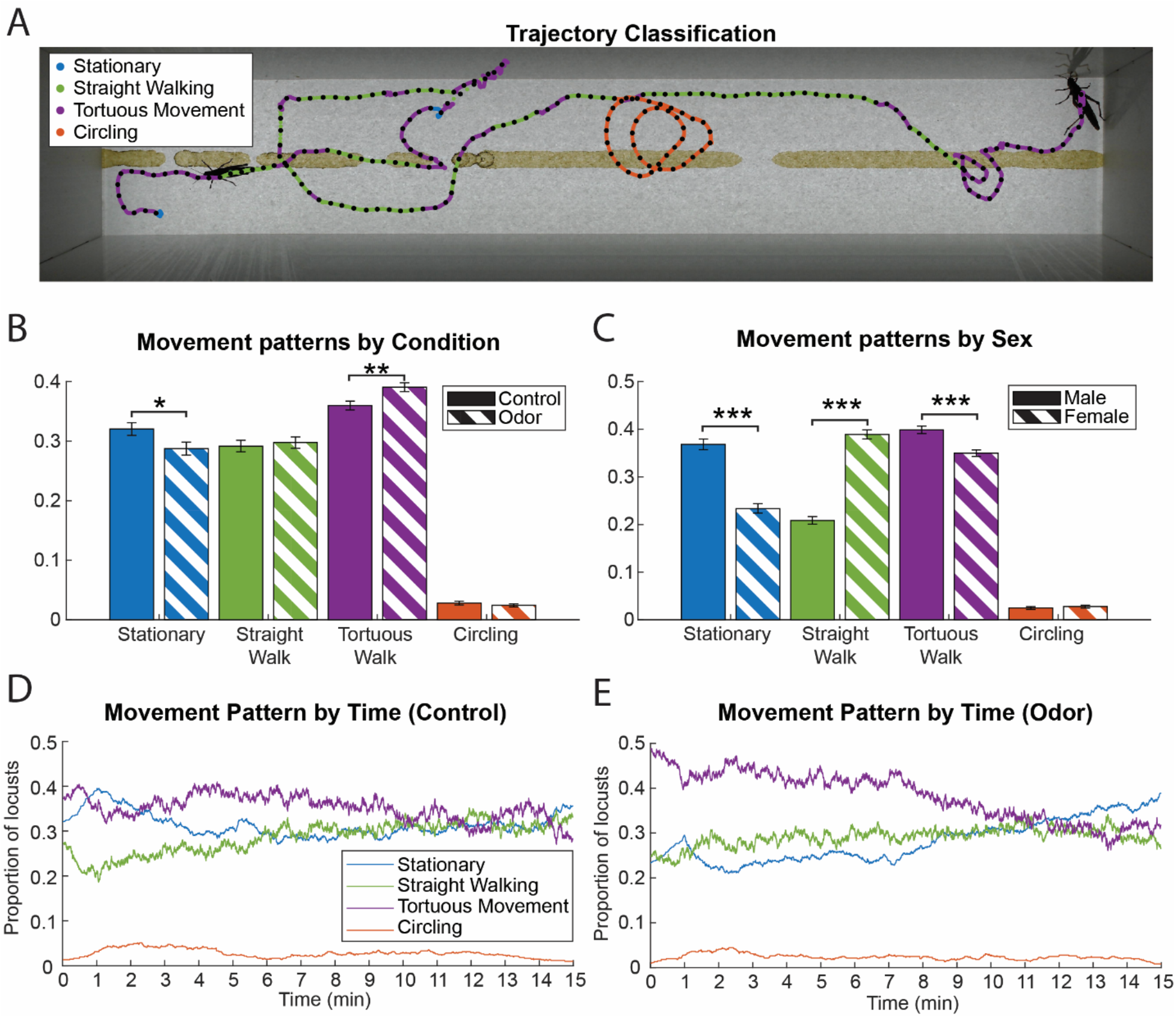
Locust walking motifs. **A)** An example walking trajectory of a locust is shown. The walking patterns with the trajectory are segmented into one of the four categories or ‘motifs’: stationary, straight walking, tortuous walking, and circling. The part of the trajectory that the locust exhibited a particular walking motif is color coded for verification. **B)** Quantification of motif expression between the odor (striped) and control (solid) conditions (n = 619). Under the odor condition, locusts spent significantly less time stationary, and significantly more time engaged in tortuous walking (* indicates p < 0.05, ** indicates p < 0.01, Wilcoxon Rank-Sum test). As can be seen in **Figure 2**, tortuous walking under visible conditions often corresponds to greater exploration of a region of the arena, suggesting that the presence of an odor spurs more investigation. (Error bars indicate s.e.m.) **C)** Quantification of motif expression between male (solid) and female (striped) locusts (male n = 323, female n = 296). Males were significantly more likely to be stationary or to engage in tortuous walking, while females were significantly more likely to engage in straight walking (*** indicates p < 0.001, Wilcoxon Rank-Sum test). Straight walking most often corresponds to walking along the edge, while interactions with the trail often are characterized by tortuous walking. (Error bars indicate s.e.m.) **D)** Motif expression as a function of time for the control condition (n=619 locusts). The X-axis indicates time and the Y-axis indicates the fraction of locusts exhibiting any one specific motif at a given point in time. Note that the fraction of locusts exhibiting straight walking increases steadily over time. **E)** Motif expression as a function of time for the odor condition (n=619 locusts). Tortuous walking formed a much larger initial proportion of the motifs expressed, but had a large dropoff as the trial continued. In contrast, stationary periods and straight walking both increased as a function of time.

Segmenting by sex instead (**Fig. 5c, Supp. Fig. 3**) revealed instead that female locusts spent significantly less time stationary and in tortuous walking (p<0.001 for both conditions; two-sided Wilcoxon signed rank test), and significantly more time walking straight (p<0.001 for both conditions; two-sided Wilcoxon signed-rank test).

Next, we examined the distribution of these walking motifs as a function of time (**Fig. 5d, e**). As can be noted, in the presence of an odor trail there were more tortuous walking bouts during early time segments, with the trends between the odor and control conditions significantly different (p=0.001 20,000 bootstrapped CI of the difference between the correlations with time excludes zero, see Methods). The amount of time spent tortuous walking decreased as the time progressed (control Spearman’s rho = -0.93, odor Spearman’s rho = -0.51, p<0.001 for both), while locusts increased the amount of straight walking (control Spearman’s rho = 0.86, odor Spearman’s rho = 0.63, p<0.001). The trends of stationary periods were also highly significant between the odor and control conditions (p=0.001 20,000 bootstrapped CI of the difference between the correlations with time excludes zero, see Methods), while straight movement and circling behavior trends were not.

The walking motifs and their dynamic transitions for each locust can also be visualized as a color-coded heatmap (**Fig. 6a, b**). The upper 323 rows are male locusts, while the remainder of the rows are female, separated by a yellow line. The largest observable difference in composition is the shift of tortuous walking (purple) among the males to straight walking (green) among the females, and the greater amount of stationary periods in the males. Furthermore, the shift from female straight walking in the control condition to a higher degree of tortuous walking in the odor condition is also visible (more purple in **Fig. 6b** bottom half compared to **Fig. 6a**). Finally, the greater complexity of behavior under the odor condition is visible as the increased speckling of behavior categories. This can be quantified by comparing bout lengths, with males (p<0.001) but not females (p=0.083) showing significant differences in bout lengths between the odor and control condition. This further indicates the greater complexity of male locust movements when presented with an odor compared to females.

**Figure 6:**
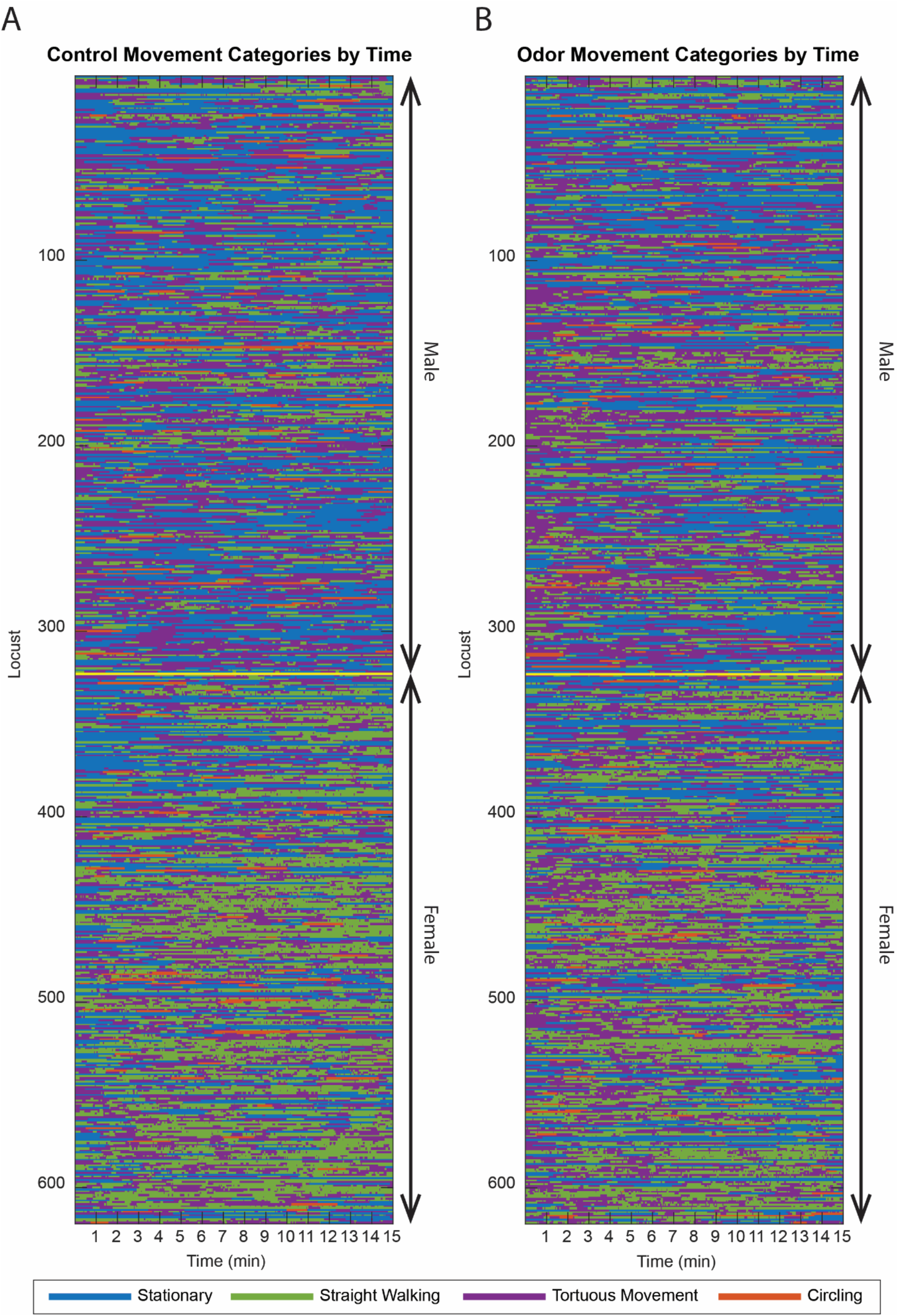
A barcode of locust movement categories. **A)** A barcode of movement categories over time for every locust during the control trial is shown (n = 619 locusts). Each row reveals the locomotory categories of each locust over the course of the trial. All male locusts are shown in the upper part of this movement matrix, and female locusts’ movement patterns are shown in the lower part of the movement matrix. The yellow line divides the male from the female locusts in the dataset. The prevalence of tortuous movement among males and straight walking among females is visible in the shift from purple in the male section to green in the female section. **B)** Similar movement category matrix for the same set of male and female locusts but for the odor trial (n = 619 locusts).

### A Markov model of odor-driven trail following behavior in locusts

We used a Markov model to summarize the behavioral data and understand the transition between the different walking. In this model, each state would correspond to one of the four motifs observed (stationary, straight walking, tortuous, circling). Time was discretized in 12.7 s bins (the median bout length across all locusts) and transition probabilities were calculated between the different walking motifs (see Methods for additional details). We fit separate models for modeling control trials vs. odor trails, male vs. female locusts, and attractive vs. repulsive odorants.

We found that the Markov models fit for different subsets of the data were highly similar (overall state transition matrix shown in **Supp. Fig. 4**). In general, each state was fairly ‘sticky,’ with a low probability of transitioning to another state. The largest transition probabilities were between straight and tortuous walking, with the second largest being between tortuous walking and circling. Stationary periods consistently had the lowest probability of transitioning to a new behavior across all conditions.

To understand the differences between different conditions, we took the difference between the transition probabilities (**Fig. 7a, b**). Between odor and control conditions, for instance, the control condition was 1.7% less likely to transition from straight walking to tortuous walking, and 2.1% more likely to remain straight walking. The comparative strength of these changes relative to the others is likely reflective of a greater tendency to walk straight up the treadmill when in the control condition, and a greater tendency to explore in tortuous movement under the odor condition.

**Figure 7:**
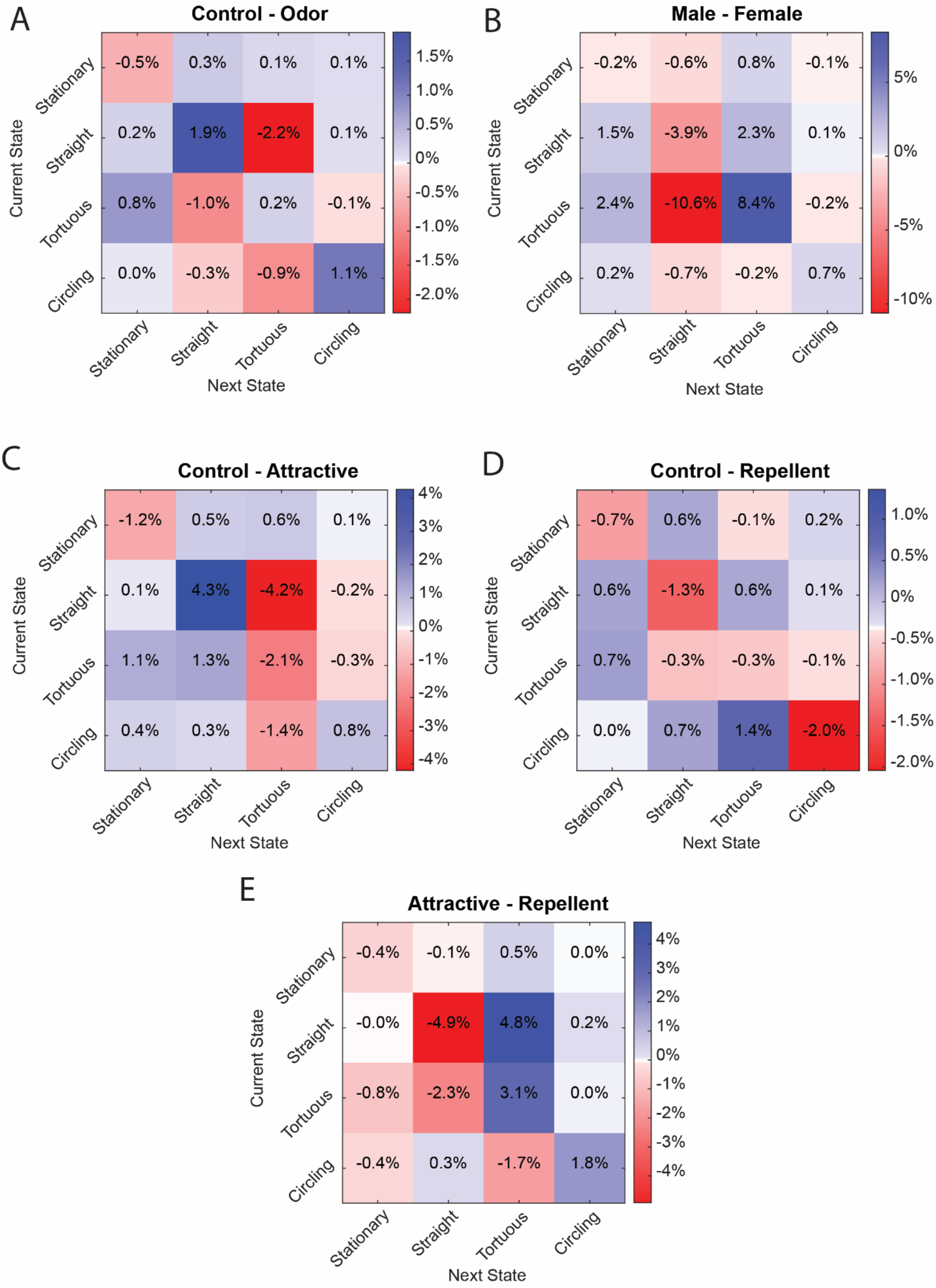
Markov models of locust walking and difference in Markov transition matrices across conditions. **A)** Differences between the transition probabilities between Markov models of locust walking in the control and odor conditions (n = 619 locusts). As expected based on overall expression levels of the walking motifs, control had a higher probability of remaining in straight walking and a reduced probability of transitioning to tortuous walking compared to the odor condition. **B**) Differences between the transition probabilities between Markov models of walking in male and female locusts (male n = 323 locusts, female n = 296 locusts). Similar to the odor and control transition probability differences, the largest difference between the sexes is the greater probability of males to continue tortuous walking, whereas females were much more likely to transition to straight walking. **C)** Differences between the transition probabilities between Markov models of walking in the control condition and the four odorants with the highest TFI (Trail following index; Isoamyl acetate 1%, Geraniol 10%, Geraniol 1%, and L-Carvone 1%, total n = 130 odor trails). Compared to the control condition, locusts exploring attractive odors are much more likely to engage in tortuous walking, typically corresponding to navigating/exploring/investigating a given length of trail more. **D)** Differences between the transition probabilities between Markov models of walking in the control condition and the four odorants with the lowest TFI odors (Citral 10%, Methyl Jasmonate 1%, Cis-3-Nonan-1-ol 1%, and Phenethylamine 1%, total n = 95 odor trails). Compared to control conditions, locusts exploring repulsive odors are less likely to transition to tortuous walking, and more likely to continue straight walking. **E)** Differences between the transition probabilities between Markov models of walking while exposed to attractive (total n = 130 odor trails) and repellant odors (total n = 95 odor trails). Locusts in attractive conditions are much more likely to transition to or continue tortuous walking, and less likely to do so for repellant odorants. However, the opposite trend holds true for circling behavior, albeit less strongly.

The differences between males and females are larger than those between the odor and control conditions (**Fig. 7a, b**). As can be expected, the largest differences in transitions occurred between straight and tortuous walking (**Fig. 7b)** with females over 10% more likely to transition from tortuous walking to straight walking than males, and over 8% less likely to continue tortuous walking. Interestingly, males are also almost 4% more likely to change from straight walking to different behavior, with 2.3% of those changes going to tortuous walking.

The differences in Markov transition probabilities between the most attractive and repulsive odorants with respect to the control behavioral trails, and with respect to each other are shown in **Fig. 7c-e**. Comparing the differences between control and attractive odorants, locusts were over 4% more likely to transition to tortuous walking from straight walking, and correspondingly less likely to continue straight walking or transition to straight walking from tortuous walking. Repellant odors, in contrast, showed a 1.3% greater likelihood to continue straight walking. Interestingly the largest difference between the repellant odor conditions and the control was a 2.0% higher probability to continue circling behavior. Finally, comparing attractive and repellant conditions, during attractive odor trail trials locusts were almost 5% more likely to transition to tortuous walking from straight walking, and over 3% more likely to continue tortuous walking. These suggest greater exploration when attractive odors are present, and a greater tendency to try and cover distance and potentially escape the odor when a repellant odor is present.

In sum, our results provide a thorough exploration of the walking behavior of locusts while following an odor trail, and how this behavior is dynamically modulated in a time, odor identity, and sex-dependent manner.

## Discussion

Locusts are a historically important model organism, with multiple unique features that make them highly useful for both neural and behavioral studies. For practical neural recording purposes, locusts have highly accessible neural systems, with straightforward access to their large antennal lobes, central complex^35,36^, and other higher brain regions, as well as motor neural processing centers in the thoracic ganglia ^37–39^. Locusts have been used to study olfactory^40–43^, visual^44^, social^45–47^, odor-sampling^48^, and foraging^49^ behavior. However, it is yet to be shown that they are able to take advantage of surface-bound odor trails for navigation, which seems particularly relevant given their known swarming/aggregation behavior. In this paper, we introduced an automated trail-following assay, and using it we were able to demonstrate that locusts are indeed able to follow odor trails. Trail-following was observed under chemosensory-only (IR) and multimodal (visual) conditions. Furthermore, we were able to examine the modulation of locomotory behavior at coarse (preference index) and fine resolution (locomotory motif expression). The trail-following behavior varied as a function of time, and as a function of odorant identity that was used to form the trail. In addition to odorant-modulated changes in activity, we also found large and significant differences in the locomotory and exploratory patterns of male and female locusts. Together, these results establish locusts as a viable model for studying odor-based navigation, introduce a new tool for doing so, and enable comparison with strategies adopted by other organisms.

Previous studies on surface-bound trial-following in ants^50^, spiders^51^, rats^13^, and dogs^1,2^ have established some commonalities in navigation strategies that are shared with airborne-plume tracking. This includes a tendency to engage in sinusoidal sweeps across the trail, called “casting” for airborne plumes when the trail is lost^13,15,50,52^. This was also observed when humans were given a surface-bound odor trail to follow ^53^. We found that locusts also engaged in sinusoidal sweeps (**Fig. 2b**). However, the zig-zagging across odor trails was predominantly observed under visible conditions and a much simpler tracking strategy appeared to be employed under IR illumination. Overall, our results provide further evidence to suggest that some strategies are shared across different organisms.

Compared to odor plume tracking there were notable differences in strategy that were also observed while following surface-bound odor trails. For animals that smell using respiration, they are able to disturb the surface by sniffing, increasing the amount of scent available for tracking^22–24^. Similarly, animals with antennae are able to probe the ground in front of them^50^, mechanically disturbing the surface while sampling the trail. As observed with ants^50^, we noted that locusts pat the surface in front of them while exploring (**Supp. Video 1**), potentially increasing the concentration of odorants available to sample. Alternately, previous studies in *Drosophila* and moths have found that wing beats were optimized to sweep air over the antennae during flight^54–56^, enabling more effective sampling of the odorants in their surroundings. However, the possession of wings for flight not only presents the opportunity for unique energetic tradeoffs but also mechanical perturbation of the surface to increase the concentration of surface-bound odorants. In fact, we noted that some locusts engaged in bouts of wing flapping without taking flight (**Supp. Video 2**); it is possible that such mechanical perturbations had the effect of increasing surface-bound odorant concentration. A systematic comparison between sampling approaches used during flight^54,55^ and walking^48,13,50^ may reveal distinctions between the strategies used and insights into mechanical^54,55^, neural^57^, and behavioral^13,50^ algorithms underlying the complex navigation that locusts and other animals engage in during daily survival.

To simplify the interpretation of how locusts moved over time, we classified locust behavior into one of four major categories including non-movement or stationary behavior and walking patterns that span the range of tortuosity (straight walk – minimum tortuosity, to circling – maximum tortuosity). We used the behavioral categorization to construct Markov models of the dynamic walking behavior observed in locusts (**Fig. 7**). In this analysis each movement category is represented as a state and the probabilities of transitions between states were computed for each insect. The observed insect behavior over time can then be summarized as a sequence or evolution of states. Such approaches have been used to successfully capture the dynamics of a number of different insect behaviors, including bee dances^58^, the foraging behavior of aphid-hunting parasitoids^59^, the stereotypy of fruit fly responses^60^, and occasionally identification of previously undescribed movement patterns^61^. Our results indicated sex-specific differences in the walking patterns of locusts. We found that the female locusts were more likely to continue walking straight whereas male locusts tend to continue zig-zagging across the odor trails. Further, the walking patterns changed depending on odorant identity and concentration. While repellant odorants elicited more straight walking, attractive odorants induced tortuous walking. These observations nicely relate patterns of avoidance and exploration with the overall valence/preference of the olfactory cue.

In sum, we have established locusts as a viable model for investigating trail following and introduced a tool for studying such behaviors. While we have used continuous trails and characterized the exploration/avoidance behavior when encountering such trails, natural odor trails tend to be fragmented and discontinuous. What local search patterns locusts would follow when encountered with those perturbations, and whether additional insights into the behavioral strategies could be gained if the locomotory responses are further fragmented into even more elementary behavioral motifs remains to be explored.

## Acknowledgments

We thank members of the Raman Lab (Washington University in St. Louis) for feedback on the manuscript. We thank Pearl Olsen and Matt Kwiatkowski for insect care. This research was supported by NSF (2021795) and ONR (N00014-19-1-2049, N00014-21-1-2343) grants to B.R.

## Author contributions

MT and BR conceived the study and designed the experiments/analyses. MT designed and constructed the treadmill, performed all the experiments, and analyzed the data. MT and BR wrote the paper. MT supervised all aspects of the work.

---

**Supp. Fig. 1:**
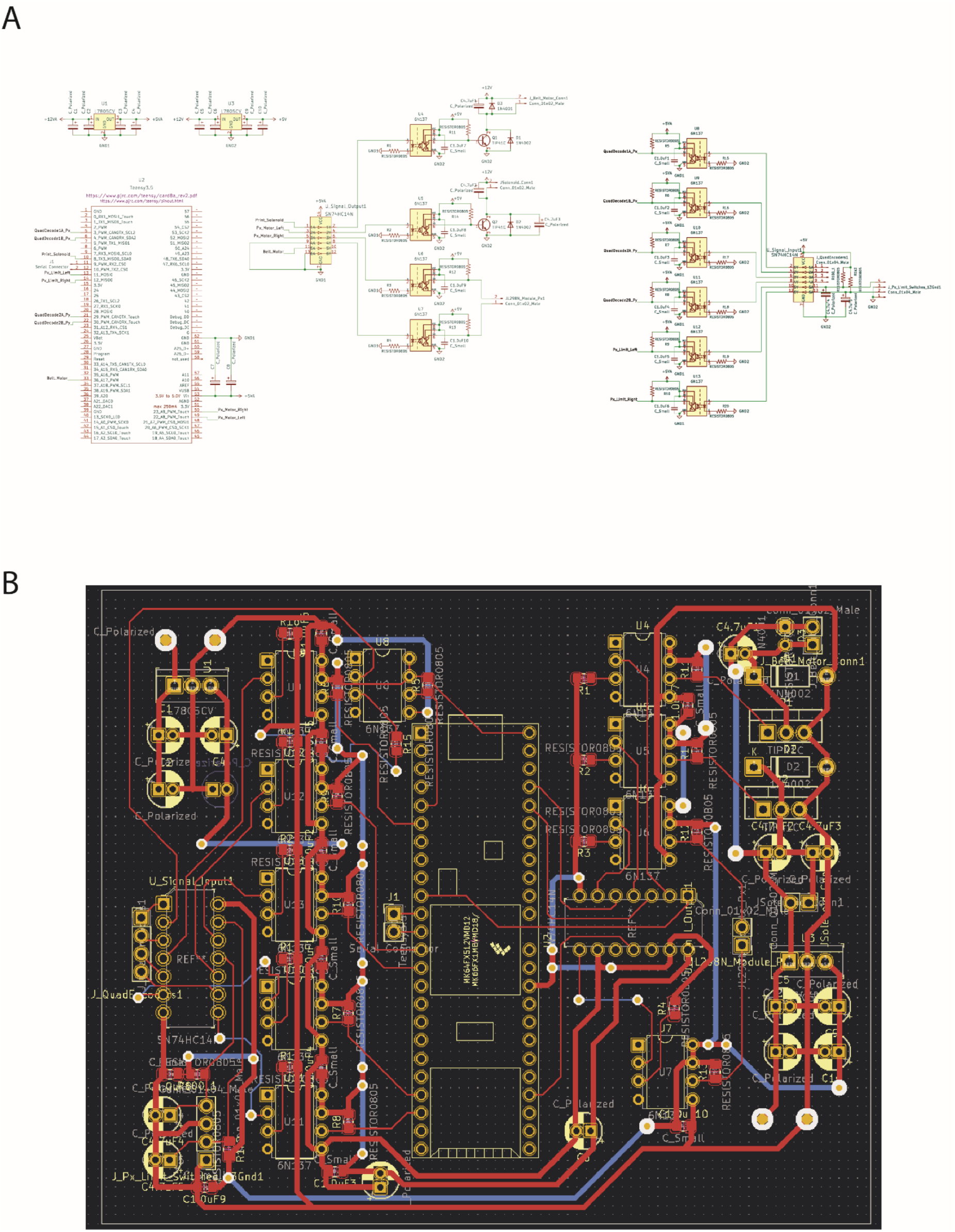
Treadmill Circuit Diagram and PCB. **A)** Connection diagram of VR Treadmill PCBN. **B)** VR Treadmill PCB schematic.

**Supp. Fig. 2:**
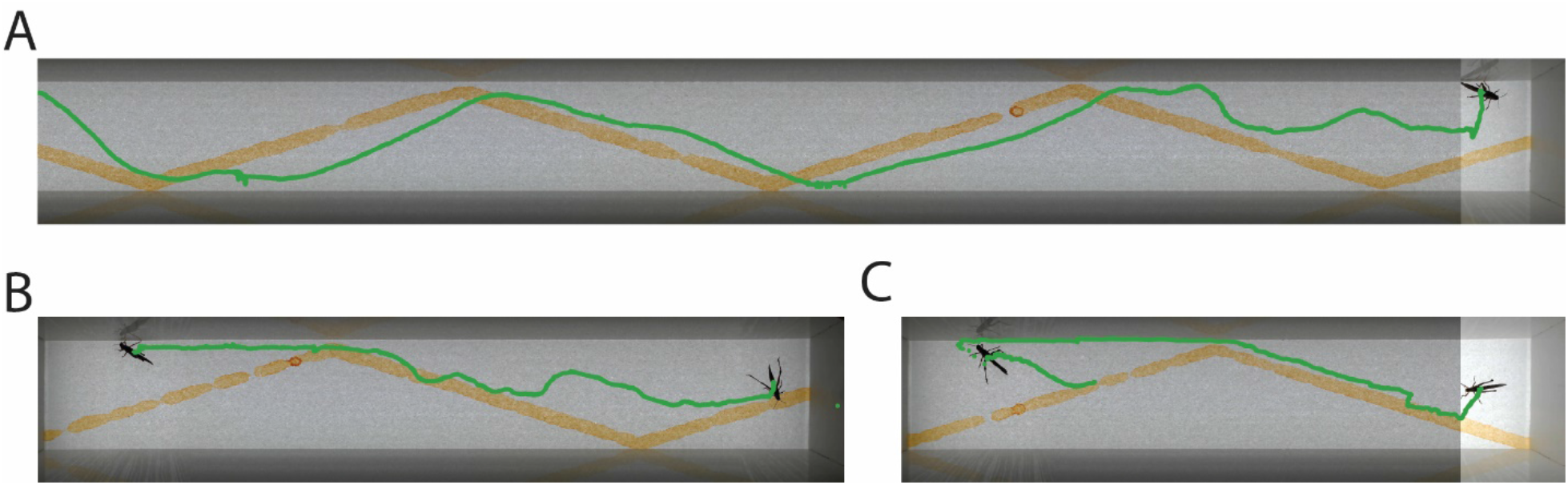
Locusts engage in trail following on complex trails. **A)** Example trail of locust following zig-zag pattern of Geraniol 10%. The trajectory is indicated by the green line. Locusts often followed on the downhill side of the trail. **B)** Same as **panel A**, however in this example, the locust can be seen to cross the trail, then return to the downhill side. However, when it encounters the wall the locust chooses to stay alongside the wall rather than continue to follow the trail. **C)** Similar example to **panel B**, however after leaving the trail, the locust later returns and starts to follow the trail again.

**Supp. Fig. 3:**
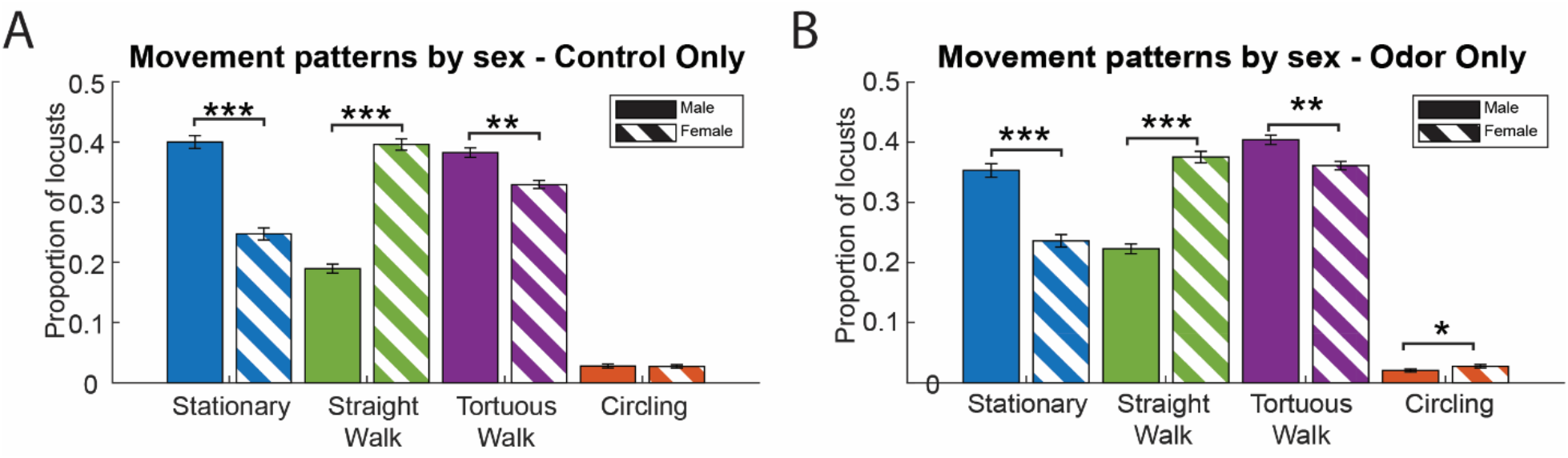
Movement category expression of male and female locusts divided by odor v. control conditions. **A)** Movement category expression of male and female locusts under the control condition (male n = 323 locusts, female n = 296 locusts). All conditions except circling are significantly different between males and females (** indicates p < 0.01, *** indicates p < 0.001, Wilcoxon Rank-Sum test. Error bars indicate s.e.m.). **B)** Movement category expression of male and female locusts under the control condition (male n = 323 locusts, female n = 296 locusts). All conditions are significantly different between males and females (* indicates p < 0.05, ** indicates p < 0.01, *** indicates p < 0.001, Wilcoxon Rank-Sum test. Error bars indicate s.e.m.).

**Supp. Fig. 4:**
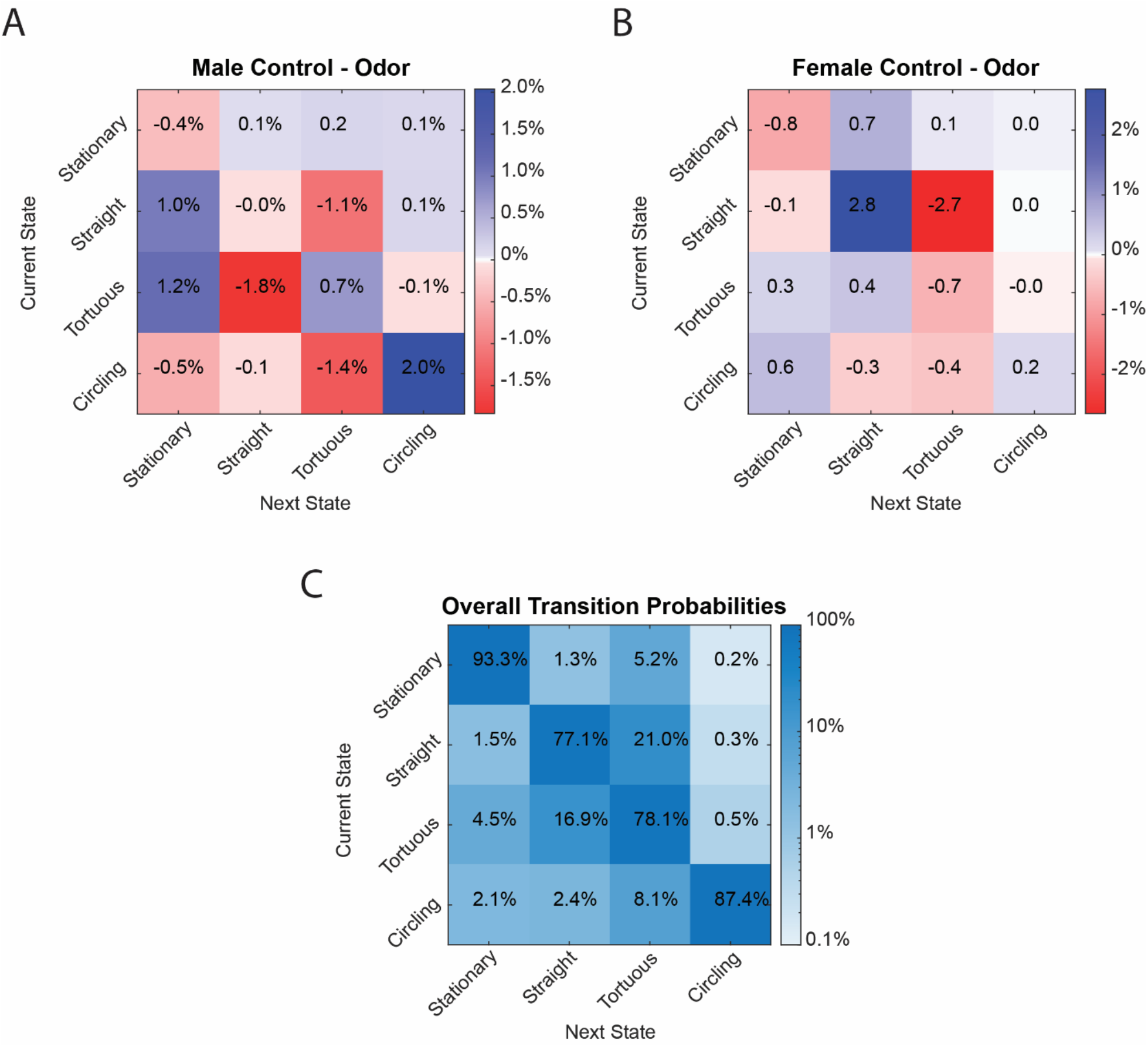
Markov Model transition probability matrices. **A)** Differences between the transition probabilities between Markov models of male locust walking in the control and odor conditions (n = 323 locusts); males have an increased likelihood to transition to circling behavior under the control condition, and are more likely to engage in straight and tortuous movement under the odor condition. **B)** Similar to **Panel A** but for female locusts (n = 296 locusts). Females have a single large shift in transition probabilities between conditions; under the control condition, they are more likely to continue straight walking, while under the odor condition, they are more likely to transition to tortuous walking. **C)** Overall transition probabilities of the Markov model across both sexes and odorant conditions (n = 619 locusts). The large values in the diagonal indicate that locusts are most likely to continue their current behavioral state than to transition to a new behavioral state, especially when stationary.

**Supp. Fig. 5:**
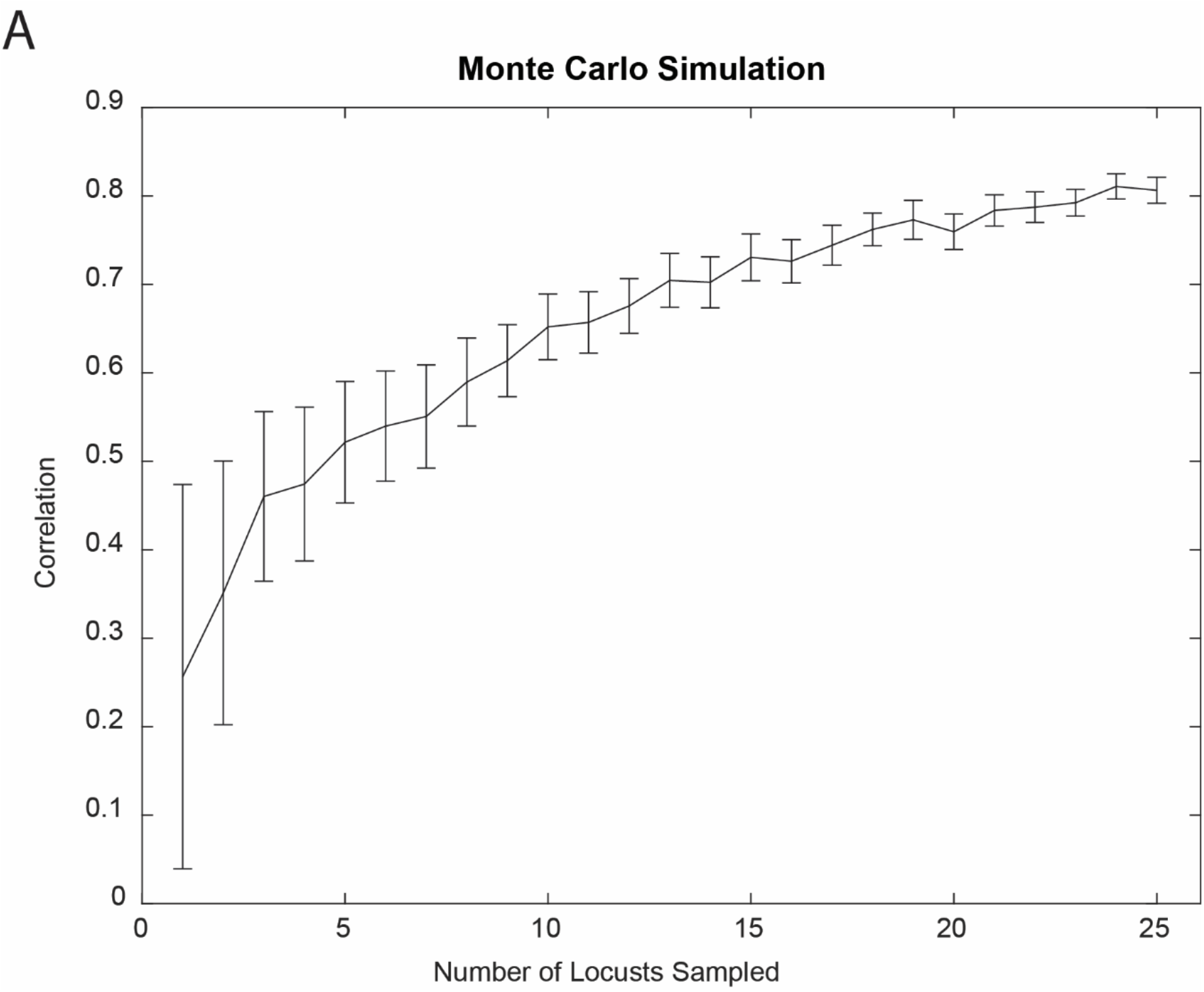
Monte Carlo simulation results. **A)** Monte Carlo simulations with replacement from each odorant sample were performed, sampling a random subset from each odorant with replacement. 100 simulations were performed for each odorant, and compared to the overall values using all samples using a correlation. A mean R^2^ value greater than 0.7 was obtained for simulations with n > 15 locusts.

